# Structural basis for the association of PLEKHA7 with membrane-embedded phosphatidylinositol lipids

**DOI:** 10.1101/2020.11.25.387084

**Authors:** Alexander E. Aleshin, Yong Yao, Amer Iftikhar, Andrey A. Bobkov, Jinghua Yu, Gregory Cadwell, Michael G. Klein, Chuqiao Dong, Laurie A. Bankston, Robert C. Liddington, Wonpil Im, Garth Powis, Francesca M. Marassi

## Abstract

PLEKHA7 (pleckstrin homology domain containing family A member 7) plays key roles in intracellular signaling, cytoskeletal organization and cell adhesion, and is associated with multiple human cancers. The interactions of its pleckstrin homology (PH) domain with membrane phosphatidyl-inositol-phosphate (PIP) lipids, are critical for proper cellular localization and function, and their inhibition is an attractive target for anti-cancer therapy. While structural data can provide insights in this area, little is known about the way in which PLEKHA7 and other PH domains interact with membrane-embedded PIPs. Here we report atomic-resolution structures of the PLEHA7 PH domain and describe the molecular mechanism for its recognition of membrane-bound PIPs. Using X-ray crystallography, nuclear magnetic resonance (NMR), molecular dynamics (MD) simulations, and isothermal titration calorimetry (ITC), we show – in atomic-level detail – that the interaction of PLEKHA7 with PIPs is multivalent and induces PIP clustering. The PIP binding mechanism is distinct from a discrete one-to-one interaction. Our findings reveal a central role of the membrane assembly in mediating protein-PIP association and provide a roadmap for the design of PLEKHA7-PIP inhibitors.

---

PLEKHA7 (pleckstrin homology domain containing family A member 7)^4^ is a major component of the cytoplasmic region of epithelial adherens junctions that functions to ensure cell-cell adhesion and tight junction integrity through its interactions with cytoskeleton proteins (Meng et al., 2008; Paschoud et al., 2014; Pulimeno et al., 2010; Rouaud et al., 2020). Increased levels of human PLEKHA7 are associated with hypertension (Levy et al., 2009) and glaucoma (Awadalla et al., 2013), and PLEKHA7 protein staining has been reported in several human cancers, including advanced breast, renal and ovarian cancer (Kourtidis et al., 2015; Tille et al., 2015), with highest occurrences in colon cancer (Castellana et al., 2012). A recent study (Nair-Menon et al., 2020) showed that disruption of the apical junction localization of RNA interference machinery proteins correlates with loss of PLEKHA7 in human colon tumors and poorly differentiated colon cancer cell lines, while restoration of PLEKHA7 expression restores proper localization of RNA interference components and suppresses cancer cell growth in vitro and in vivo. Moreover, we have observed (Jeung et al., 2011) that PLEKHA7 associates with proteins of the membrane-bound KRAS signaling nanocluster in colon cancer cells, and that *plekha7* gene knock down inhibits the proliferation of colon cancer cells with mutated KRAS, but not normal cells with wild-type KRAS. PLEKHA7, therefore, represents an attractive druggable target for the selective inhibition of signaling and its tumor-related functions in colorectal cancer. Despite its importance, the molecular basis for the role of PLEKHA7 in these disorders is poorly understood.

PLEKHA7 is one of nine family members characterized by a 120-residue N-terminal pleckstrin homology (PH) domain. PH domains are found in more than 100 different proteins, they are known for their ability to bind phosphatidyl-inositol-phosphate (PIP) lipids within cell membranes, and their association with PIPs is essential for intracellular signaling, cytoskeletal organization and the regulation of intracellular membrane transport (DiNitto and Lambright, 2006; Lemmon, 2007). Notably, PH domains can be selectively targeted by small molecules that inhibit their signaling function (Indarte et al., 2019; Meuillet et al., 2003; Meuillet et al., 2010). In the case of PLEKHA7, the PH domain is required for establishing proper subcellular localization through its interactions with PIPs (Wythe et al., 2011), and thus, could offer a new avenue for attacking cancer by inhibiting PLEKHA7 localization.

Following the first NMR structures of pleckstrin (Yoon et al., 1994) and β-spectrin (Macias et al., 1994), the structures of many other PH domains have been reported, both in their free state or complexed with soluble inositol phosphate (IP) small molecules (Moravcevic et al., 2012). These have provided important functional insights, but little is still known about the way in which PH domains associate with full-length PIPs incorporated in membranes, and nothing is known about the structural basis for PIP binding by PLEKHA7. Using a multidisciplinary approach that combined X-ray crystallography, NMR, all-atom MD simulations, and ITC binding measurements, we have determined the structures of the PLEHA7 PH domain and characterized its interactions with membrane-embedded PIPs with atomic-level detail. The data reveal three PLEKHA7 binding sites for PIP, and demonstrate that confinement of PIP molecules by the membrane assembly is essential for promoting the multivalent association of one PH domain with multiple PIPs leading to membrane surface localization.

## Results and discussion

### Structure of the PLEKHA7 PH domain

We obtained crystals for two states of the PH domain of PLEKHA7 – one ligand-free (PHA7_APO_) and the other bound to sulfate from the crystallization buffer (PHA7_S_) – and we determined two structures, which refined to 2.80 Å and 1.45 Å resolution, respectively (Fig. 1A, B; Fig. S1A-C; Table S1). Each structure crystallized in a different space group and has two copies of the protein per crystallographic asymmetric unit, but the protein-protein interfaces differ in each case, and do not appear to reflect biologically functional dimers. PLEKHA7 adopts the PH domain signature fold (Lemmon, 2007; Lietzke et al., 2000): a seven-stranded β-barrel, capped at one end by a 16-residue α-helix. The barrel opening is highly positively charged and lined by the β1-β2, β3-β4 and β6-β7 loops which are hypervariable in both length and sequence across the PH domain family. The K173-K183-R185 motif, at the end of β1 and start of β2, forms the canonical IP binding site of PH domains (Moravcevic et al., 2012). In the case of PHA7_S_, this site – denoted here as site I – is occupied by one of two sulfate anions resolved in the structure. No electron density was observed for either the β6-β7 loop or C-terminal residues S285-R298, which were included in the sequence of PHA7_S_ (residues 164-298) but deleted in PHA7_APO_ (residues 164-285). The structures show that the PH domain is confined to residues 164-285, and subsequent NMR, ITC and MD simulation studies utilized this trimmed sequence, henceforth referred to as PHA7 (Fig. S1A).

**Fig. 1.**
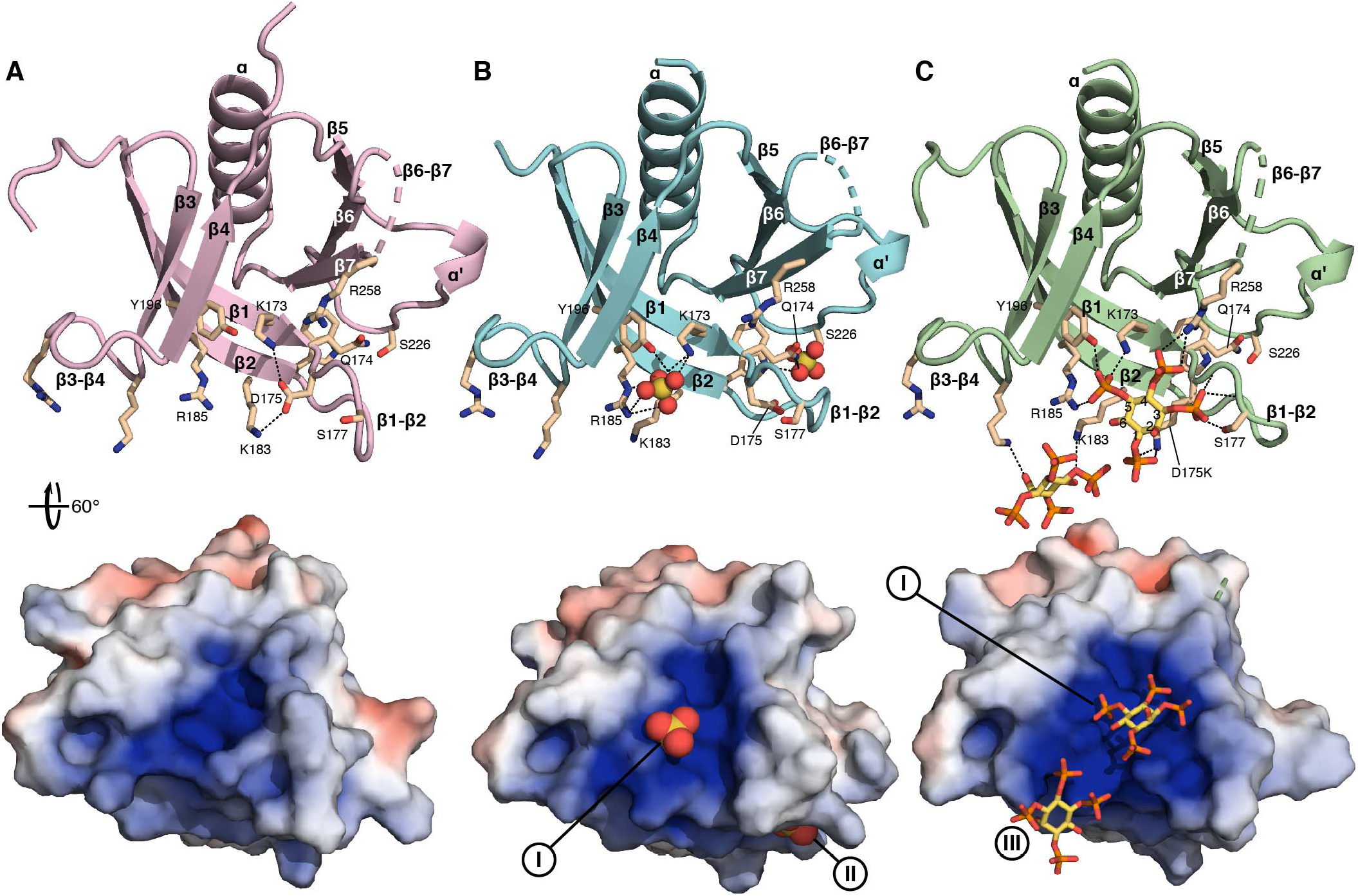
Structure of the PLEKHA7 PHD. **(A)** PHA7_APO_. **(B)** PHA7_S_ bound to sulfate (yellow/orange spheres). **(C)** Mutant PHA7-D175K bound to soluble IP(3,4,5)P_3_ (yellow/orange sticks). Key residues are shown as sticks. Dashes denote protein-protein, protein-sulfate or protein-IP(3,4,5)P_3_ polar contacts (<4Å). The 60° surface representations are colored by electrostatic potential from −5 kT/e (red) to +5 kT/e (blue). The positions of the three binding sites (I-III) for sulfate or phosphate are marked.

To further examine the protein conformation and dynamics in solution, we prepared ^15^N/^13^C labeled PHA7 for NMR experiments. The assigned NMR chemical shifts (^1^HN, ^15^N, ^13^CA and ^13^CB) show that the protein adopts the same structure in solution as in its crystalline form, and that the long β6-β7 loop forms a random coil (Fig. S2A). The ^1^H/^15^N nuclear Overhauser effect (NOE) data show that the β-strands, and helices (α1, α’) all have similar (>0.8) NOE intensities, consistent with uniform backbone dynamics and conformational order (Fig. S2B). By contrast, the termini and loops have distinctly lower NOE intensities, and hence greater extents of dynamics and lower conformational order. High flexibility is observed for β6-β7 where the intensities approach zero, and also for β1-β2 and β3-β4, loops that are important for defining the recognition of specific PIP phosphorylation states by other PH domains (Lemmon, 2007).

PLEKHA7 shares 50-75% amino acid sequence identity with the other PLEKHA family members (Fig. S3A), but it is unique in containing a 20-residue insertion between β6 and β7. Long β6-β7 loops are found in other members of the larger PH domain superfamily. In GRP1 and ARNO (Cronin et al., 2004; Lietzke et al., 2000), the β6-β7 loop forms a β-hairpin that elongates the β-barrel, generating a more extensive pocket for binding IP, while in the case of ANLN, the loop does not appear to be involved in IP binding. The β6-β7 sequence of PHA7 differs from those of GRP1, ARNO and ANLN.

### Interactions with soluble IP heagroups and membrane-embedded PIPs

To assess the affinity of PHA7 for membrane-embedded PIP lipids, we performed co-sedimentation experiments with PIP-containing liposomes. While PHA7 does not sediment with pure dipalmitoyl-phosphatidyl-choline (diC16-PC) liposomes, incubation with liposomes containing 10% molar diC16-PI(3,4,5)P_3_ resulted in abundant co-sedimentation (Fig. 2A). Notwithstanding this evident high affinity for PIP-rich membranes, our attempts to co-crystallize PHA7 with soluble IP headgroups – IP(4,5)P_2_, IP(3,4)P_2_ or IP(3,4,5)P_3_ – resulted in the apo form of the protein, without bound ligand, regardless of crystallization or soaking conditions. This is in line with the μM-range dissociation constants (K_d_) that we measured by ITC for the interaction of PHA7 with soluble IPs (Fig. 2B; Fig. S4; Table 1). The combined data indicate that PHA7 has modest affinity for soluble IPs but appreciably higher affinity for PIP lipids incorporated in membranes. A similar effect has been reported for the PH domain of kindlin 3 whose binding affinity for PIP was enhanced 1,000 fold upon PIP incorporation in nanodiscs (Ni et al., 2017).

**Fig. 2.**
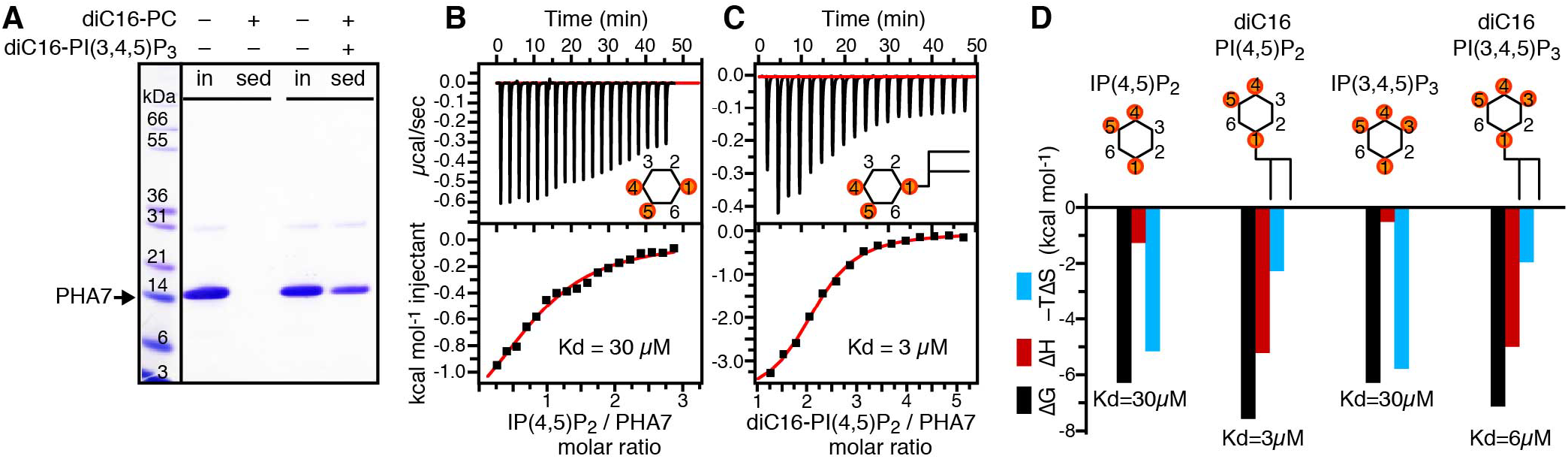
Interactions of PHA7 with PIP lipids and soluble IPs. **(A)** SDS-PAGE analysis of PHA7 co-sedimentation with diC16-PC liposomes prepared with or without 10% molar diC16-PI(3,4,5)P_3_. Monomeric PHA7 (arrow) migrates with apparent molecular weight of ~15 kDa. **(B-D)** Representative ITC binding isotherms (B, C) and free energies (D) measured for titrations of PHA7 with soluble IP(4,5)P_2_ and IP(3,4,5)P_3_, or nanodiscs containing 10% molar diC16-PI(4,5)P_2_ and diC16-PI(3,4,5)P_3_. Continuous red lines are the best fits of the data to a single-site binding model, used to extract the values of the dissociation constant (Kd).

**Table 1.**
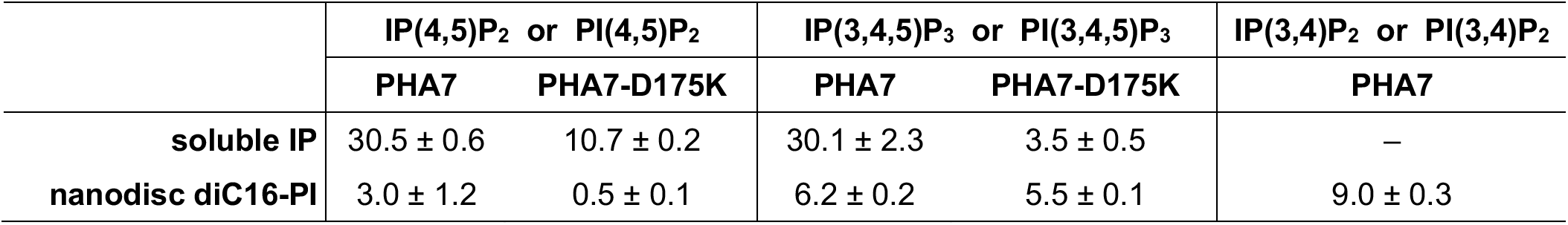
ITC values of Kd (μM) for the interactions of wild-type PHA7 or mutant PHA7-D175K with soluble IP headgroups or diC16-phosphatidyl inositol (diC16-PI) incorporated in lipid nanodiscs.

|  | IP(4,5)P<sub>2</sub> or PI(4,5)P<sub>2</sub> |  | IP(3,4,5)P<sub>3</sub> or PI(3,4,5)P<sub>3</sub> |  | IP(3,4)P<sub>2</sub> or PI(3,4)P<sub>2</sub> |
| --- | --- | --- | --- | --- | --- |
|  | PHA7 | PHA7-D175K | PHA7 | PHA7-D175K | PHA7 |
| **soluble IP** | 30.5 ± 0.6 | 10.7 ± 0.2 | 30.1 ± 2.3 | 3.5 ± 0.5 | – |
| **nanodisc diC16-PI** | 3.0 ± 1.2 | 0.5 ± 0.1 | 6.2 ± 0.2 | 5.5 ± 0.1 | 9.0 ± 0.3 |

To analyze this effect quantitatively, we performed ITC experiments with PHA7 and PIP-enriched lipid nanodiscs – nanometer-size discoidal membranes that are stabilized by two copies of a membrane scaffold protein (MSP) derived from the apolipoprotein ApoA1 (Bayburt et al., 2002). We prepared nanodiscs with either pure diC16-PC or a 1/9 molar mixture of diC16-PIP and diC16-PC. Remarkably, the ITC binding affinity measured by titrating the protein with PIP-nanodiscs is enhanced by more than one order of magnitude relative to soluble IP headgroups (Fig. 2C; Fig. S4; Table 1), in line with the co-sedimentation results. Moreover, PHA7 now appears to display some – albeit very modest – selectivity (p<0.05) for PI(4,5)P_2_ > PI(3,4,5)P_3_ > PI(3,4)P_2_ lipid membranes.

The enhanced affinity for membrane-incorporated PIPs suggests that additional factors – beyond a pure one-to-one PHA7-IP interaction – play a role in recruiting the protein to the membrane surface. The binding free energies derived from ITC offer some insights in this regard (Fig. 2D). The association of PHA7 with free IPs is highly entropy driven, with binding thermodynamics characterized by very small values of favorable enthalpy (ΔH) and a much greater entropic component (ΔS). The situation is reversed for the association of PHA7 with membrane-embedded PIPs, where the enthalpic component dominates. This effect may reflect a reduction of PIP conformational flexibility imposed by the membrane scaffold, as well as the interaction of one PHA7 molecule with multiple PIP molecules incorporated in the nanodisc membrane. The additional free energy observed for membrane-integrated PIPs may be attributed to a change in the probability of binding that results from confinement of multiple PIPs by the membrane, according to the principles of binding free energy additivity (Jencks, 1981). Such superadditivity is well known in drug development, where chemical linking of weakly-binding fragments results in molecules with binding free energy greater than the sum of the individual fragments. In the case of PHA7-PIP, membrane confinement replaces chemical linking to yield enhanced affinity.

### Structural basis for binding soluble IPs

In the absence of co-crystals, the structure of PHA7_S_ provides useful insights about the protein’s interactions with the IP moiety. In PHA7_S_ (Fig. 1B; Fig. S5A) one sulfate anion occupies site I, where it is coordinated by the side chains of K173 in β1, R185 in β2 and Y196 at the end of β3. A second sulfate anion binds on the opposite side of the β1-β2 loop – denoted here as site II – where it is coordinated by the side chains of Q174 and W182, which define the start and the end of the β1-β2 loop, and S226 in the middle of the β5-β6 loop. While IP binding at this second site is considered “atypical”, and thought to occur only in PH domains that lack the canonical binding site (Moravcevic et al., 2012), it has been observed experimentally for the PH domains of spectrin (Hyvonen et al., 1995), Tiam1 and ArhGAP9 (Ceccarelli et al., 2007), and ASAP1 (Jian et al., 2015), where both canonical and atypical sites are simultaneously bound to short-chain PI(4,5)P_2_.

Notably, while the structures of PHA7_APO_ and PHA7_S_ are superimposable, with root mean square deviation (rmsd) of 0.49/0.52 Å (chains A-A/B-B) for CA atoms in the barrel core, they differ appreciably in the β1-β2 loop where the rmsd jumps to 2.54/2.42 Å (chains A-A/B-B) (Fig. S5B). In the apo structure, the β1-β2 loop folds towards the barrel opening and adopts a β-turn conformation that is restrained by a network of hydrogen bonds involving the K173 and K183 amino groups, the S177 hydroxyl, and the D175 carboxylate oxygens (Fig. 1A; Fig. S5C, D). The D175 side chain points into binding site I, at the center of a clasp-like structure formed by the K173 and K183 side chains. The temperature factors of the β1-β2 loop are twice greater than for the rest of the protein, and the loop has a distinct conformation in each of the two molecules of the crystal structure, reflecting its flexibility (Fig. S1B). In the sulfate-bound structure, on the other hand, the β1-β2 loop folds away from the barrel opening (Fig. 1B; Fig. S5C, D). The D175 side chain is disengaged from the K173-K183 clasp: It points into the loop, forming hydrogen bonds with its main chain atoms, and away from the sulfate anion bound in site I. The β1-β2 temperature factors, while higher than those of the barrel proper, are lower than those observed in PHA7_APO_, and the loop has the same conformation in both molecules of the structure, indicating that the bound sulfate stabilizes the β1-β2 loop conformation (Fig. S1C).

The structures point to D175 and the β1-β2 loop as important mediators of the interaction of PHA7 with IP headgroups. A previous study (Carpten et al., 2007) identified a “sentry” Glu residue in the PH domain of AKT positioned to disfavor binding of a phosphate group at the 3-position of the inositol ring and define a preference for IP(4,5)P_2_ over IP(3,4)P_2_ and IP(3,4,5)P_3_. Changing this sentry Glu to Lys, an AKT mutation that occurs in many tumors, reverses the preference and increases the affinity for IPs by ~10-fold.

To examine the role of D175 in PHA7, we generated the D175K mutant and characterized its structure and IP binding properties. ITC measurement show that PHA7-D175K has a three-fold greater affinity for IP(4,5)P_2_ (K_d_=10.7 μM) and nine-fold greater affinity for IP(3,4,5)P_3_ (K_d_=3.5 μM) compared to wild-type (Fig. S4; Table 1). Notably, PHA7-D175K co-crystallized with soluble IP(3,4,5)P_3_. The structure refined to a resolution of 2.43 Å (Fig. 1C, Table S1) with two copies of the protein per crystallographic asymmetric unit. Once again, no electron density was observed for the long β6-β7 loop, indicating that it remains mobile and disordered, without apparently contributing to IP binding.

The two molecules of PHA7-D175K within the crystallographic asymmetric unit coordinate two IP(3,4,5)P_3_ moieties sandwiched between them: One at the site I, and the second associated more peripherally with yet another electropositive patch – denoted here as site III – formed by the side chains of K183, R185, K198, and R201 that protrude from β2 and from the β3-β4 loop (Fig. S6). The structure reflects at least four distinct modes of IP binding by PHA7. In molecule A of the asymmetric unit the phosphate group at the fifth position of the inositol ring occupies the same position as the sulfate anion in the structure of wild-type PHA7_S_, while in molecule B the binding geometry is flipped such that it is the phosphate group in first position that coincides with the sulfate binding site (Fig. S6B, C). Binding of the peripheral IP(3,4,5)P_3_ at site III is similarly mirrored in each copy of PHA7-D175K, resulting in two possible binding geometries. Moreover, the involvement of K183 and R185 in both sites I and III reflects plasticity in the PHA7-IP interaction.

To further examine the interaction of wild-type PHA7 with IPs, we performed NMR experiments in solution. Addition of free IP(3,4,5)P_3_ into ^15^N-labeled PHA7 resulted in specific ^1^H/^15^N chemical shift perturbations that map to the PHA7 barrel opening (Fig. 3A-B). In line with the ITC data, NMR reflects a weak binding interaction with fast exchange binding dynamics (Williamson, 2013). No major structural reorganization of either the core or the β6-β7 loop of PHA7 is observed. Notably, however, signals from all three binding sites identified by crystallography are perturbed (Fig. 3D, E), with the largest changes observed for the β1-β2 and β3-β4 loops and residues at the start of β7, including key residues (Q174, D175, S177, K183, R185, Y260) identified in the structures of PHA7_S_ and PHA7-D175K. We conclude that the interaction of PHA7 with soluble IP headgroups is structurally plastic, with three potential binding sites for inositol phosphate localized across the electropositive barrel opening, and low selectivity for a specific orientation of the IP(3,4,5)P_3_ moiety.

**Fig. 3.**
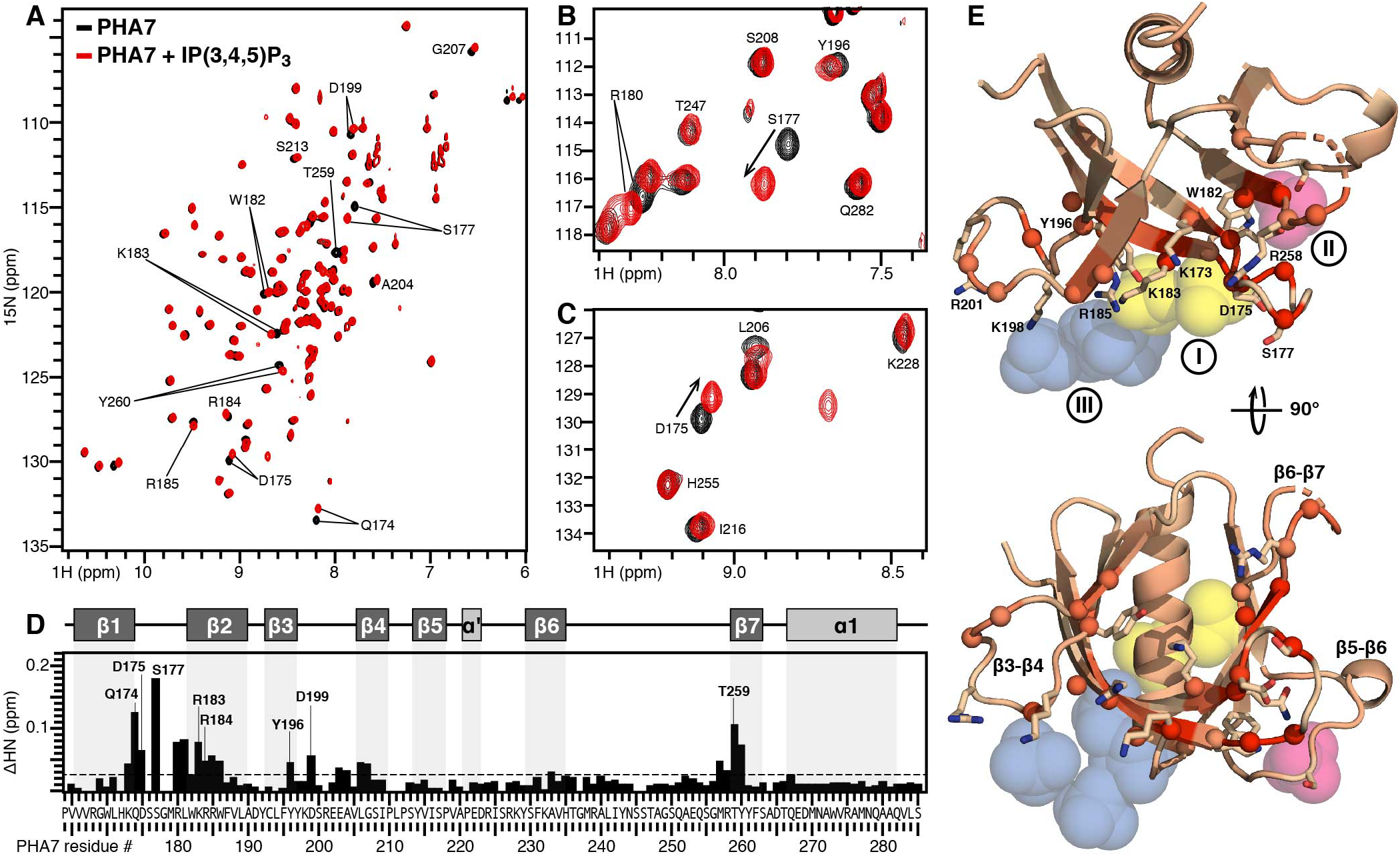
Mapping the interaction of PHA7 with soluble IP(3,4,5)P_3_. **(A-C)** ^1^H/^15^N HSQC NMR spectra of ^15^N labeled wild-type PHA7 domain acquired with (red) or without (black) 1.5 molar equivalents of soluble IP(3,4,5)P_3_. The spectra were acquired at 15°C. Selected regions of the spectra are expanded (B) to highlight specific perturbation sites. **(D)** Profile of ^1^H/^15^N chemical shift perturbations induced by IP(3,4,5)P_3_ across the sequence of PHA7. Bars represent the combined difference (ΔHN) of amide ^1^H and ^15^N chemical shifts. The protein secondary structure is outlined at the top. **(E)** Orthogonal views of the structure of PHA7_APO_. Colors reflect the magnitude of ΔHN from 0 ppm (wheat) to the maximum value (red). Highly perturbed sites (> 1 standard deviation of the values of ΔHN) have CA atoms shown as spheres. Key side chains of three IP binding sites (I-III) are shown as sticks. The positions of sulfate or phosphate groups identified in the structures of PHA7_S_ and PHA7-D175K are shown as spheres, superimposed on the structural model. They denote binding sites I (yellow), II, (pink) and III (blue).

### Structural basis for binding membrane PIPs

The structural and ITC binding data show that PHA7 has modest affinity for freely soluble IP ligands but appreciably higher affinity for PIP-rich lipid bilayers, pointing to the importance of the PIP membrane environment. To explore the interaction with PIP membranes at the atomic level, we performed NMR experiments with ^15^N labeled PHA7 and PIP-nanodiscs. Incubation of PHA7 with diC16-PC nanodiscs, resulted in no detectable NMR spectral changes (Fig. 4A; S7A). This is consistent with the liposome co-sedimentation results and confirms the inability of PHA7 to bind membranes devoid of PIP. Incubation with nanodiscs containing 10% molar diC16-PI(3,4,5)P_3_ gave a dramatically different result: The NMR spectrum was effectively obliterated, except for peaks from sites in the termini and the flexible β6-β7 loop (Fig. 4B; S7B). We interpret such massive peak suppression to reflect immobilization of the relatively small, 125-residue protein, caused by its association with the comparatively large nanodisc membrane. Moreover, since both PC and PIP nanodisc preparations have similar and homogeneous size distributions (~8 nm radius; Fig. S8), we attribute the peak suppression effect to a PIP-dependent association of the protein with the nanodisc membrane that is long-lived on the msec time scale of ^1^H/^15^N correlation NMR experiment.

**Fig. 4.**
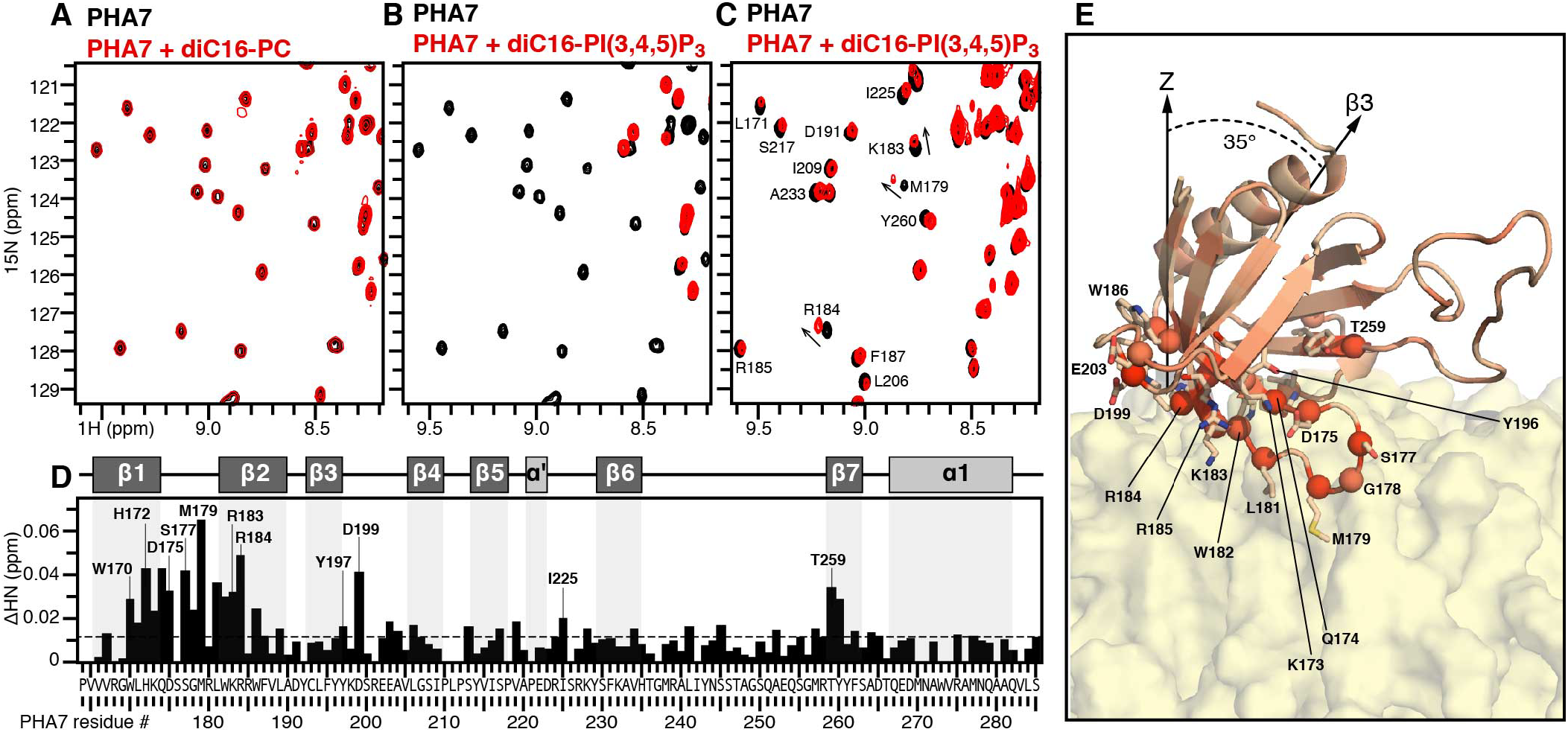
Mapping the interaction of PLEKHA7 with diC16-PI(3,4,5)P_3_ nanodiscs. **(A-C)** ^1^H/^15^N HSQC NMR spectra of ^15^N-labeled wild-type PHA7 obtained before (black) or after (red) incubation with 1.5 molar equivalents of lipid nanodiscs prepared with 100% diC16-PC (A), or 9/1 molar diC16-PC and diC16-PI(3,4,5)P_3_ (B, C). The spectra were acquired at 15°C (A, B) or 45°C (C). **(D)** Profile of ^1^H/^15^N chemical shift perturbations induced by the association of PHA7 with diC16-PI(3,4,5)P_3_ nanodiscs, at 45°C, across the protein sequence. **(E)** Snapshot (taken at 890 ns) of a MD simulation of PHA7 with a diC16-PI(3,4,5)P_3_-rich lipid bilayer membrane. PHA7 colors reflect the magnitude of NMR chemical shift perturbation from 0 ppm (wheat) to the maximum value (red). Highly perturbed sites (> 1 standard deviation of the values of ΔHN) have CA atoms shown as spheres and side chains shown as sticks. The membrane is represented as a molecular surface.

Increasing the temperature restored the NMR spectrum (Fig. 4C; Fig. S7C), and while this is expected to enhance the binding exchange dynamics, it also allowed us to map specific chemical shift perturbations that reflect the interaction of PHA7 with the PIP nanodisc membrane. The perturbations (Fig. 4D) mirror the profile observed with free IP(3,4,5)P_3_ (Fig. 3C), but map to more extended regions of β1 and β2, and include sites in β5-β6. The largest perturbations map to the β1-β2 hairpin, the β3-β4 loop, and the start of β7, implicating binding sites I, II and III in PIP-mediated membrane association, and indicating that PHA7 uses all three to dock onto the PIP-membrane surface.

All atom MD simulations provide molecular context for the NMR data. We performed ten independent, unrestrained, 1-μs MD simulations of PHA7 for each of three different lipid bilayer membranes similar to the experimental nanodiscs (Table S2). All ten simulations with either PI(4,5)P_2_ or PI(3,4,5)P_3_ resulted in rapid membrane surface association of PHA7, which remained membrane-bound throughout the course of 1-μs simulation, while little evidence of binding to PC-only membranes was observed (Fig. S9). In all cases, PHA7 adopts a preferred orientation at the PIP membrane surface, with the β3 strand at an average angle of 35° from the membrane normal, the C-terminal helix exposed to bulk water, and sites I, II and III docked on the membrane (Fig. 4E; Fig. S10).

Notably, the interaction patterns from unrestrained MD simulations mirror the experimental NMR perturbation profile (Fig. 5A, B). Protein sites with NMR signals that are sensitive to PIP-nanodiscs also interact with the PIP membrane surface in the MD simulations. Viewed in a snapshot of MD simulation with a PI(3,4,5)P_3_ membrane (Fig. 4E), the NMR perturbations are associated with membrane-binding sites. These include conserved hydrophobic residues (L181, M179) in the β1-β2 loop that insert deeply into the hydrophobic core of the membrane, thereby providing added stability to the interaction. The PH domain orientation at the membrane surface is remarkably similar to that observed from EPR-guided MD simulations of GPR1 with PIP membranes (Lai et al., 2013). In that study, hydrophobic interactions between GPR1 side chains and the membrane core were also observed, indicating that superficial membrane penetration helps stabilize a specific geometry of PH domain association with the membrane surface.

**Fig. 5.**
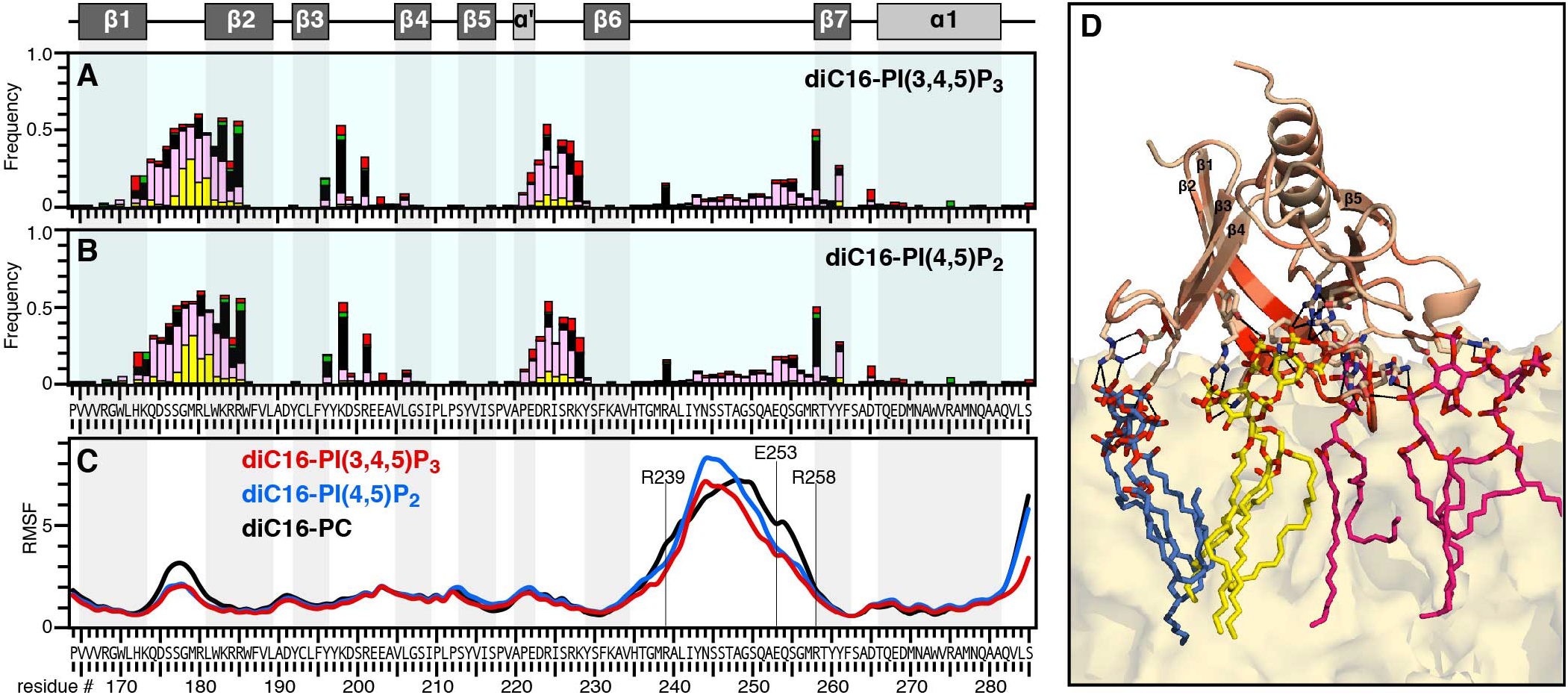
Structure and dynamics of the PLEKHA7 PH domain. **(A, B)** MD interaction profile of PHA7 with diC16-PI(3,4,5)P_3_ or diC16-PI(4,5)P_2_ membranes. The bars represent frequency of occurrence within 4 Å of water (light blue), Na^+^ (red) or Cl^−^ (green) ions, phospholipid tails (yellow), PC headgroups (pink), or PIP headgroups (black). Each data point is the average of ten independent MD simulations over the last 500 ns of 1-μs MD trajectories. **(C)** Time-averaged RMSF calculated for PHA7 heavy atoms over the last 500 ns of 1-μs MD trajectories for ten independent simulations in diC16-PI(3,4,5)P_3_ (red), diC16-PI(4,5)P_2_ (blue), or diC16-PC (black) membranes. **(D)** Snapshot (taken at 890 ns) of MD simulation of PHA7 with a diC16-PI(3,4,5)P_3_ membrane. PHA7 colors reflect the magnitude of NMR chemical shift perturbation from 0 ppm (wheat) to the maximum value (red). Key side chains with close (< 4 Å) contacts to PIP headgroups are shown as sticks. The membrane is represented as a molecular surface (yellow). Colors of bound PIP molecules (lines) represent their association with binding sites I (yellow), II (pink), or III (blue).

In line with this observation, the time-averaged root mean-squared fluctuations (RMSF), calculated for heavy atoms over the last 500 ns of 1-μs MD trajectories, show that the β1-β2 loop experiences a marked reduction in conformational flexibility upon binding PIP membranes (Fig. 5C). By contrast this effect is not observed for PC membranes. Moreover, while the long β6-β7 loop remains highly flexible in both the soluble and membrane-associated states of PHA7, the width of its flexible region is significantly reduced. At the molecular level, the loop fluctuations are dampened at its edges, where R239 and R258 engage with a PIP phosphate group and E253 helps stabilize the interaction.

Finally, closer examination of the PHA7-PIP membrane assembly reveals multivalent interactions of the protein with PIPs. A total of seven PI(3,4,5)P_3_ molecules establish close contacts (> 4Å) with the protein: Two PIP molecules associate with each of sites I and III, and three associate in the periphery of site II (Fig. 5D; Fig. S11). In this way, PHA7 recruits a cluster of PIP molecules to its periphery. The data demonstrate that PHA7 established multivalent interactions with membrane PIPs. These are primarily electrostatic, involving basic sidechains and inositol ring phosphate groups, but also include hydrogen bonds from polar side chains, and hydrophobic contacts between nonpolar side chains and the lipid acyl chains. These results explain the enhanced affinity of PHA7 for PIP membranes compared to freely soluble IPs.

## Conclusions

PLEKHA7 relies on the interactions of its PH domain with PIPs for proper localization to the plasma membrane surface and for its normal functions (Wythe et al., 2011). Using a multidisciplinary approach, we have defined the major elements of PIP recognition by the PLEKHA7 PH domain and demonstrated the important role of the membrane scaffold in mediating its interaction with PIPs.

Although PH domains share a common fold and sequence homology, their PIP affinities and selectivities vary broadly. ITC shows that PLEKHA7 has modest affinity for free IP headgroups and no detectable selectivity for phosphorylation at specific inositol ring sites. By contrast, the binding affinity for PIP membranes is highly enhanced. Compared to freely soluble IP headgroups, the interaction with membrane PIPs is enthalpy-driven and exhibits slight selectivity for PI(4,5)P_2_ (> PI(3,4,5)P_3_ > PI(3,4)P_2_) relative to the other phosphorylation types. PI(4,5)P_2_ is the major PIP lipid present in the inner leaflet of the plasma membrane of mammalian cells. Its phosphorylation by PI 3-kinase (PI3K) produces PI(3,4,5)P_3_, a key second messenger in pathways related to cell survival and metabolism, while dephosphorylation of PI(3,4,5)P_3_ by phosphatases produces PI(3,4)P_2_ (Cantley, 2002). Low PIP selectivity is in line with the observations (Wythe et al., 2011) that PLEKHA7 membrane targeting is not linked to the PI3 kinase-dependent conversion of PI(4,5)P_2_ to PI(3,4,5)P_3_ in cells, and that its PH domain is promiscuous for PI(4)PP_1_, PI(4,5)P_2_, PI(3,4,5)P_3_, as well as other non-inositol phospholipids with accessible phosphate groups, in lipid overlay assays.

The crystal structures reveal three positively charged binding sites for the phosphate groups of the PIP headgroup moiety, and the NMR data show that all three sites engage with soluble IP headgroups, forming an extended binding surface at the open end of PH domain β-barrel. Notably, NMR studies with PIP nanodiscs show that the same binding sites, plus conserved hydrophobic residues in the β1-β2 loop, engage with membrane-embedded PIPs leading to membrane surface association of the PH domain that is long-lived on the msec time scale. Moreover, the MD simulations show that PLEKHA7 interacts with the PIP membrane in two ways: by engaging the phosphate groups of multiple PIP molecules through electrostatic interactions with its three binding sites, and by inserting hydrophobic side chains into the hydrophobic core of the lipid bilayer membrane. This multivalent binding interaction results in long-lived association of PH domain with the PIP membrane, on the μs time scale of the MD simulations.

Our results indicate that PLEKHA7 induces membrane PIP clustering. We note that our simplified nanodiscs and membrane systems have both restricted geometries and PIP concentrations that are ten times higher than cells – conditions that are likely to favor cluster formation. In cells, the reverse situation, where PLEKHA7 binds to pre-formed PIP clusters, is also possible. While the cellular levels of PI(4,5)P_2_ are low (~1% of total lipid), clustering is thought to increase its local concentration at specific membrane sites. A recent study (Wen et al., 2018) demonstrated that physiological levels of divalent and trivalent metal ions can induce cluster formation of very low PIP concentrations (< 0.05 mol% of total lipids). The ability to form such cation-bridged PIP clusters at extremely low concentrations reveals an important property of this central lipid and provides evidence for the formation of distinct pools of PIP in cellular membranes, with fundamental consequences for biological function. As noted (Wen et al., 2018), cation-induced PIP clustering would dampen the functions of proteins that bind free PIP lipids while enhancing the functions of proteins that bind clusters preferentially.

Clustering of PIP molecules around PH domains has also been observed in coarse-grained MD simulations, and has been proposed to contribute to the binding free energy (Yamamoto et al., 2020). Notably, two cooperative PIP-binding sites have been observed experimentally in the PH domain of ASAP1, and this has been proposed to enable rapid switching between active and inactive states during cellular signaling (Jian et al., 2015). For PLEKHA7, we identified three binding sites, and found that all three are engaged upon addition of either soluble IPs or PIP membranes, leading us to conclude that multivalent association is operative in both settings. Nevertheless, the membrane assembly is fundamentally important. PLEKHA7 establishes additional hydrophobic interactions with the membrane hydrocarbon core and adopts a preferred orientation at the membrane surface. Moreover, confinement of PIP molecules by the membrane scaffold reduces the degrees of freedom of the system to a single binding interface. We propose that these factors cooperate to lower the binding free energy and enhance the affinity of protein association with PIP membranes.

While most experimental studies have focused on the interactions of PH domains with soluble IP headgroups, our results provide atomic-level insights about the affinity of PLEKHA7 for full-length membrane-embedded PIPs. They highlight the central function of the membrane assembly and provide a roadmap for understanding how the functions of the PH domain integrate with the signaling, adhesion and nanoclustering functions of full-length PLEKHA7 in cells.

## Methods

### Protein preparation

Protein sequences (Fig. S1) of human PLEKHA7 were cloned into the BamHI and XhoI restriction sites of the pGEX-6P-1 plasmid, and expressed in *E. coli* BL21 cells. For crystallography and ITC the bacterial cells were grown at 37°C, in LB media. For NMR studies, the bacteria were grown in M9 minimal media containing (^15^NH_4_)_2_SO_4_ and/or ^13^C-glucose (Cambridge Isotope Laboratories), to obtain isotopically labeled protein. Protein expression was induced by adding 1 mM isopropyl-1-thio-β-D-galactopyranoside to the culture when the cell density reached OD_600_ = 0.6. After growing the cells for an additional 15 hrs at 18°C, they were harvested by centrifugation (6,000 x g, 4°C, 15 min) and stored at −80°C overnight.

Cells harvested from 2 L of culture were suspended in 35 mL of buffer A (25 mM Na/KPO_4_, pH 7.4, 200 mM NaCl, 2 mM DTT, 1 mM EDTA), supplemented with protease inhibitors (cOmplete Mini EDTA-free cocktail; Roche), and lysed using a French Press. The soluble fraction was isolated as the supernatant from centrifugation after cell lysis, and purified by passing first on Glutathione Sepharose beads (GE Healthcare), and second by cation exchange chromatography (HiTrap SP column, 5 ml, GE Healthcare) with a linear gradient of NaCl in buffer B (20 mM Tris-Cl, pH 8, 1 mM DTT, 1 mM EDTA). The protein was concentrated to 10 mg/ml and stored frozen at −80°C. Purified protein was transferred to appropriate buffers by size exclusion chromatography (Superdex 75 10/300, GE Healthcare) and stored at 4°C.

### Nanodisc preparation

Briefly, the phospholipids were dissolved in 1 mL of nanodisc buffer (20 mM Tris-Cl, pH 7.5, 100 mM NaCl, 1 mM EDTA) supplemented with Na-cholate to obtain a final 1:2 molar ratio of lipid to cholate. MSP1D1Δh5 was produced in *E. coli*, as described (Hagn et al., 2013), dissolved in 700 μL of nanodisc buffer, and combined with the lipid solution. After incubation at room temperature for 1 hr, Biobeads SM-2 (Biorad; 2 g; prewashed in nanodisc buffer) were added, and the mixture was further incubated at room temperature, with gentle mixing, for 12 hr. The Biobeads were removed by centrifugation and the resulting nanodiscs were washed twice with one sample volume of nanodisc buffer. The nanodisc solution was concentrated using a 10 kD cutoff Vivaspin concentrator (Viva Products) and replaced with NMR buffer to obtain 500 μL of 0.2 mM nanodiscs. The nanodisc concentration was estimated by measuring A280 of MSP1D1Δh5, of which there are two copies per nanodisc. The distribution of PIP molecules among nanodiscs is assumed to be homogeneous, but this may not be the case. Nanodiscs were prepared with 100% diC16-PC, or 9/1 molar mixtures of diC16-PC with diC16-PI(3,4,5)P_3_, diC16-PI(4,5)P_2_, or diC16-PI(3,4)P_2_. Analytical size exclusion chromatography (Superdex 75 10/300 GL column, GE Healthcare) was performed in NMR buffer to assess nanodisc size homogeneity (Ding et al., 2015).

### ITC experiments

ITC experiments were performed at 23°C, with all components in ITC buffer (15 mM Tris pH 7.6, 70 mM NaCl, 0.5 mM TCEP), using an iTC200 instrument (MicroCal). Titrations with soluble IP headgroups were performed with 50 μM PHA7 in the ITC cell and 0.5 mM soluble IP molecules in the injection syringe. Titrations with PIP nanodiscs were performed with 15 μM nanodiscs in the ITC cell and 0.5 mM PHA7 in the injection syringe. The protein concentrations were estimated by measuring absorbance at 280 nm (A_280_). Nanodisc concentrations reflect the A280 measurement of MSP1D1Δh5. Integrated heat data were processed and analyzed with ORIGIN software (Microcal). The data were fit to a single-site binding model to extract the values of the dissociation constant for each titration.

### Liposome co-sedimentation assays

Dry lipids, either 100% diC16-PC, or a 9/1 molar mixture of diC16-PC and diC16-PI(4,5)P_2_, were suspended in 2 mL of buffer (20 mM MES pH 6.0, 100 mM NaCl, 2 mM DTT, 1 mM EDTA) at a concentration of 3 mg/mL, then sonicated in a bath sonicator until the suspension became translucent, marking the formation of small unilamellar vesicles. A 200 uL solution of vesicles was mixed with a 20 uM solution of PLEKHA7 and incubated at room temperature for 2 h. Liposomes were then harvested by centrifugation in a BECKMAN Airfuge (A-100/30 rotor, 91,000 rpm) for 20 h. The supernatant and sediment fractions were separated and analyzed by SDS-PAGE.

### Crystallization, X-ray data acquisition and structure determination

All PHA7 crystals were obtained using the sitting drop method. For ligand-free PHA7_APO_, protein solution (13 mg/ml in 180 mM NaCl, 20 mM Tris pH 8, 50 mM BisTris pH 6.0, 0.7 mM TCEP, 6 mM Na azide) was mixed with an equivalent volume of crystallization solution (20% PEG 3350, 20 mM MgCl_2_, 20 M NiCl_2_, 100 mM HEPES pH 7) and equilibrated at room temperature. For PHA7_S_, protein solution (30 mg/ml in 180 mM NaCl, 20 mM sodium phosphate pH 6.5, 5 mM DTT) was mixed with an equivalent volume of crystallization solution (25% glycerol, 2 M (NH_4_)_2_SO_4_) and equilibrated at room temperature. Crystals were frozen without the addition of a cryoprotectant. For PHA7-D175K, 1.5 mM IP(3,4,5)P_3_ and protein solution (15 mg/ml in 180 mM NaCl, 20 mM Tris pH 8, 30 mM BisTris pH 6, 0.5 mM TCEP, 6 mM Na azide) were mixed with an equivalent volume of crystallization solution (20% PEG 3350, 200 mM Na acetate) and equilibrated at room temperature. Crystals were frozen after addition of glycerol to final concentration 20% v/v.

X-ray diffraction data for PHA7_APO_ and PHA7_S_ were collected at the Advanced Light Source (Berkeley, CA) beamline 8.3.1, with a wavelength 1.116 Å and temperature of 100 K. The data for PHA7-D175K were collected on a Rigaku diffractometer with R-axis detector, with a wavelength of 1.54 Å and temperature of 100 K. The data were processed using the CCP4 suite (Winn et al., 2011), to resolution of 2.80 Å (PHA7_APO_), 1.45 Å (PHA7_S_) and 2.43 Å (PHA7-D175K). The structure of PHA7_S_ was solved first, with molecular replacement guided by the structure of PEPP1 (PDB: 1UPR; 52% identity). Phenix.AutoBuild (Adams et al., 2010) was used for initial model building, followed by several rounds of manual model inspection and correction in Coot (Emsley et al., 2010) and refinement by phenix.refine (Adams et al., 2010) and Refmac5. The structures of PHA7_APO_ and PHA7-D175K were solved using Phaser (52) using the structure of PHA7_S_ as molecular replacement model, and refined as for PHA7_S_.

Molprobity (Chen et al., 2010) and the PDB validation server were used for structure validation throughout refinement. All structures had Ramachandran statistics with more than 95% of residues in favored positions and less then 1% outliers. Poisson-Boltzmann electrostatics were calculated in PyMOL using APBS (Baker et al., 2001). Illustrations were prepared using PyMol.

### NMR experiments

Samples for NMR studies were transferred to NMR buffer (20 mM MES pH 6.0, 100 mM NaCl, 1 mM TCEP, 1 mM EDTA). For studies of PLEKHA7 with nanodiscs, the purified PH domain was added directly to preformed nanodiscs prepared in NMR buffer. Assignments of the solution NMR resonances from N, HN, CA and CB were obtained using HNCA (Grzesiek and Bax, 1992) and HNCACB (Wittekind and Mueller, 1993) experiments. Chemical shifts were referenced to the H2O resonance (Cavanagh et al., 1996). Secondary structure was characterized by analyzing the chemical shifts with TALOS+ (Cornilescu et al., 1999; Shen et al., 2009). The total differences (ΔHN) in amide ^1^H and ^15^N chemical shifts due to the C-terminus or peptide binding were calculated by adding the changes in ^1^H (ΔH) and ^15^N (ΔN) chemical shifts using the equation ΔHN= (1/2) [(ΔH)^2^ + (ΔN/5)^2^]. The NMR data were processed and analyzed using NMRPipe (Delaglio et al., 1995), Sparky (Goddard and Kneller, 2004) and NMRView (Johnson and Blevins, 1994). The NMR pulse sequences are described in detail in the literature (Bax and Grzesiek, 1993; Cavanagh et al., 1996; Clore and Gronenborn, 1998; Ferentz and Wagner, 2000; Fesik and Zuiderweg, 1990; Kay, 2001).

### Molecular dynamics simulations

All-atom MD simulations were performed using the CHARMM36(m) force fields for protein and lipids (Brooks et al., 2009; Klauda et al., 2010), with the TIP3P water model (Jorgensen et al., 1983) in 100 mM NaCl. All systems were prepared using CHARMM-GUI *Solution Builder* and *Membrane Builder* (Jo et al., 2008; Jo et al., 2009; Lee et al., 2019), and equilibrated with the CHARMM-GUI standard protocol. The temperature and pressure were maintained at 318.15 K and 1 bar. MD production simulations were conducted with OpenMM (Eastman et al., 2013) for 1 μs, and the last 500 ns of trajectories were used for analysis. The initial structural model was taken from the crystal structure of _PHA7S_; all ligands were removed and residues 236-254 were modeled using GalaxyFill.

Ten MD simulations were performed for each of the three membrane systems (Table S3), for a total of thirty independent simulations. The initial system size of each replica was 90 Å × 90 Å × 175 Å to accommodate for all initial components, yielding a total of approximately 125,000 atoms per replica. All analyses were performed using CHARMM, and models were visualized with VMD and PyMOL.

### Data availability

The structural atomic coordinates have been deposited in the Protein Data Bank with accession codes 7KK7, 7KJO and 7KJZ. The assigned NMR chemical shifts have been deposited in the BMRB databank with accession code 50512.

## ^4^Abbreviations

IP(4,5)P_2_: inositol (1,4,5) triphosphate
IP(3,4)P_2_: inositol (1,3,4) triphosphate
IP(3,4,5)P_3_: inositol (1,3,4,5) tetraphosphate
ITC: isothermal titration calorimetry
MD: molecular dynamics
NMR: nuclear magnetic resonance
PH: pleckstrin homology
PIP: phosphatidyl inositol phosphate
PI(4,5)P_2_: phosphatidyl inositol (4,5) diphosphate
PI(3,4)P_2_: phosphatidyl inositol (3,4) diphosphate
PI(3,4,5)P_3_: phosphatidyl inositol (3,4,5) triphosphate
PLEKHA7: pleckstrin homology domain containing family A member 7
HSQC: heteronuclear single quantum coherence

## Acknowledgments

This work was supported by grants from the National Institutes of Health (GM118186, CA179087, CA160398, CA030199) and the National Science Foundation (MCB1810695).

## Declaration of interests

The authors declare that they have no conflicts of interest with the contents of this article. The content is solely the responsibility of the authors and does not necessarily represent the official views of the National Institutes of Health.

## Author contributions

F.M.M., R.C.L. and G.P. conceived and guided the project.

A.E.A. performed crystallography and determined the structures.

Y.Y and J.Y. performed NMR experiments and data analysis.

A.E.A. G. C., M.G.K. and L.A.B. prepared samples

A.A. B. performed ITC experiments and data analysis.

W.I. guided the MD simulations.

A.I. and C.D. performed and analyzed the MD simulations.

F.M.M. prepared figures and wrote the manuscript.

All authors discussed the results and contributed to manuscript preparation.

## Additional information

**Supplementary information** is available for this paper.

**Correspondence and requests for materials** should be addressed to F.M.M.

## Notes

### Competing Interest Statement

The authors have declared no competing interest.

