## Supplementary Figures and Tables for "Structural basis for the association of PLEKHA7 with membrane-embedded phosphatidylinositol lipids"

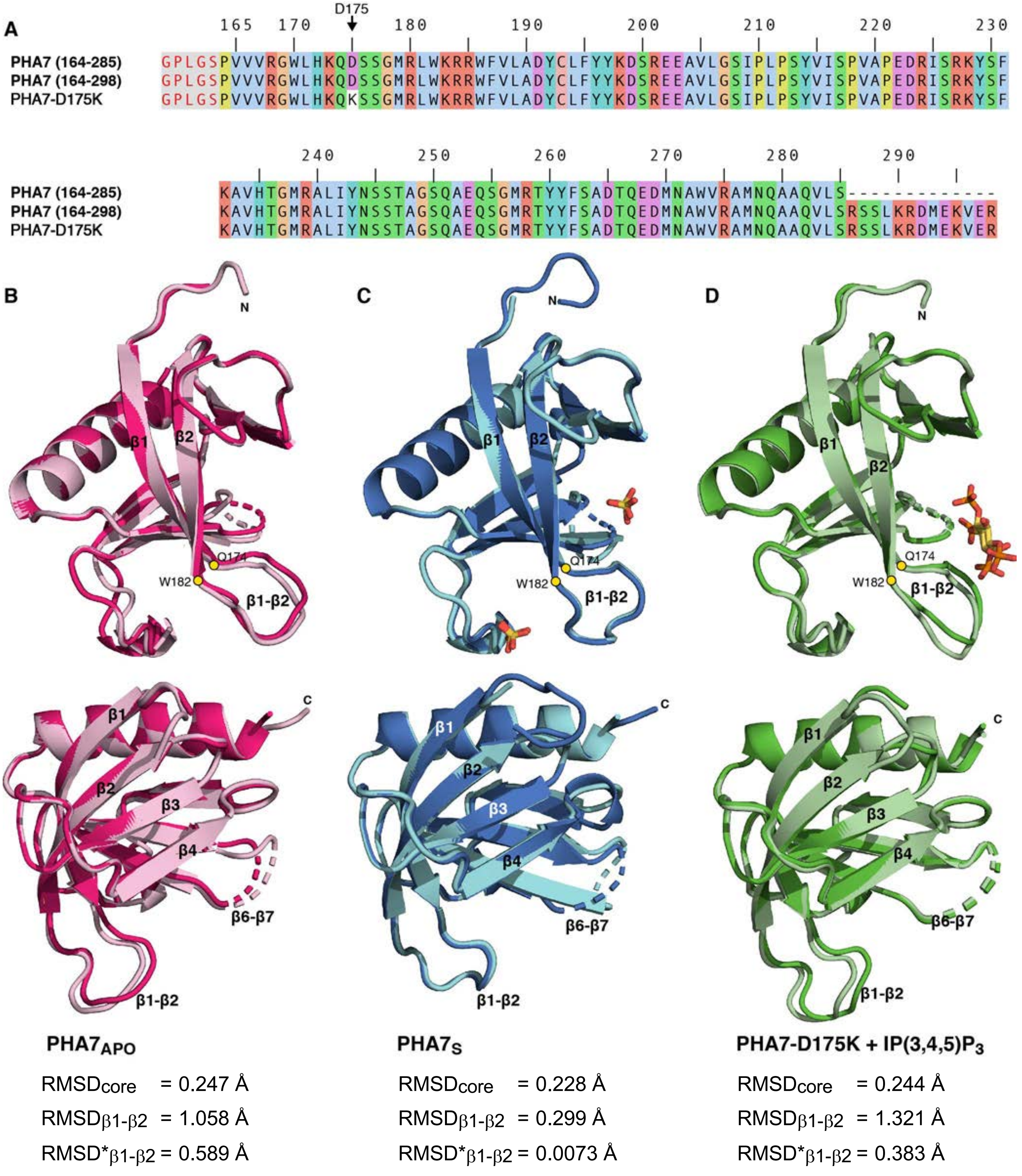


**Figure S1. Sequences and structure of PHA7. (A)** Amino acid sequences of wild-type PHA7 and mutant PHA7-D175K used for structure determination, NMR and ITC studies. The sequence begins at residue P164 after N-terminal cloning artifacts (red font over gray background). **(B-D)** X-ray crystal structures of $PHA7_{APO}$ (pink/magenta), $PHA7_{S}$ (cyan/blue) and PHA7-D175K (pale green/green). The two copies of the protein resolved for each crystallographic asymmetric unit are superimposed. $RMSD_{core}$ is calculated for the best matching CA atoms after alignment of the entire molecule. $RMSD_{\beta1\text{-}\beta2}$ is calculated for CA atoms of the β1-β2 loop (Q174-W182) after alignment of the entire molecule. $RMSD^{*}_{\beta1\text{-}\beta2}$ is calculated for CA atoms of the β1-β2 loop after alignment of the β1-β2 loop. Structures were rendered with Pymol (DeLano, 2005, *www.pymol.org*).

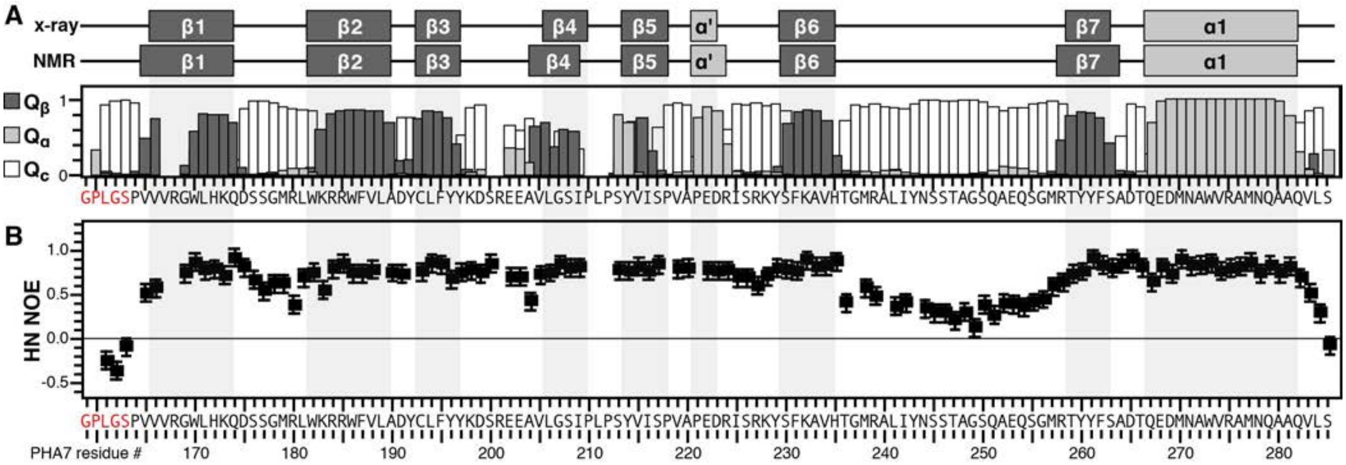


**Figure S2. Summary of the secondary structure and dynamics of PHA7 (164-285) derived from NMR. (A)** Secondary structure from TALOS-based chemical shift analysis. Bars depict the accuracy (Q) of prediction for α-helix (Q-α), β-strand (Q-β), or random coil (Q-c). Elements of the secondary structure derived from crystallography and NMR are depicted above the plot. **(B)** Heteronuclear $^{1}H/^{15}N$ NOE relative intensities for $PHA7_{APO}$.

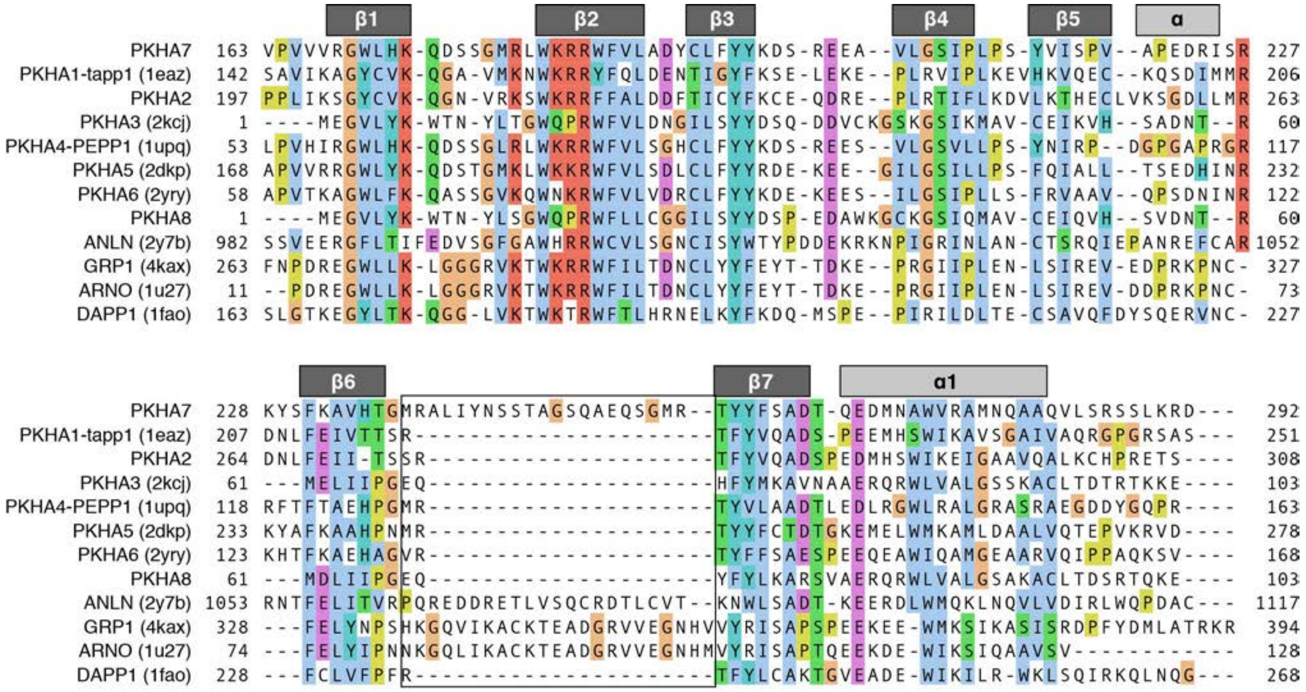


**Figure S3**. **Structure-based sequence alignment of PLEKHA family and other PH domains.** Sequence alignments were generated and colored with Clustal Omega using Jalview (Clamp et al., 2004, *Bioinformatics* 20: 426). The conserved secondary structure is shown above the sequences.

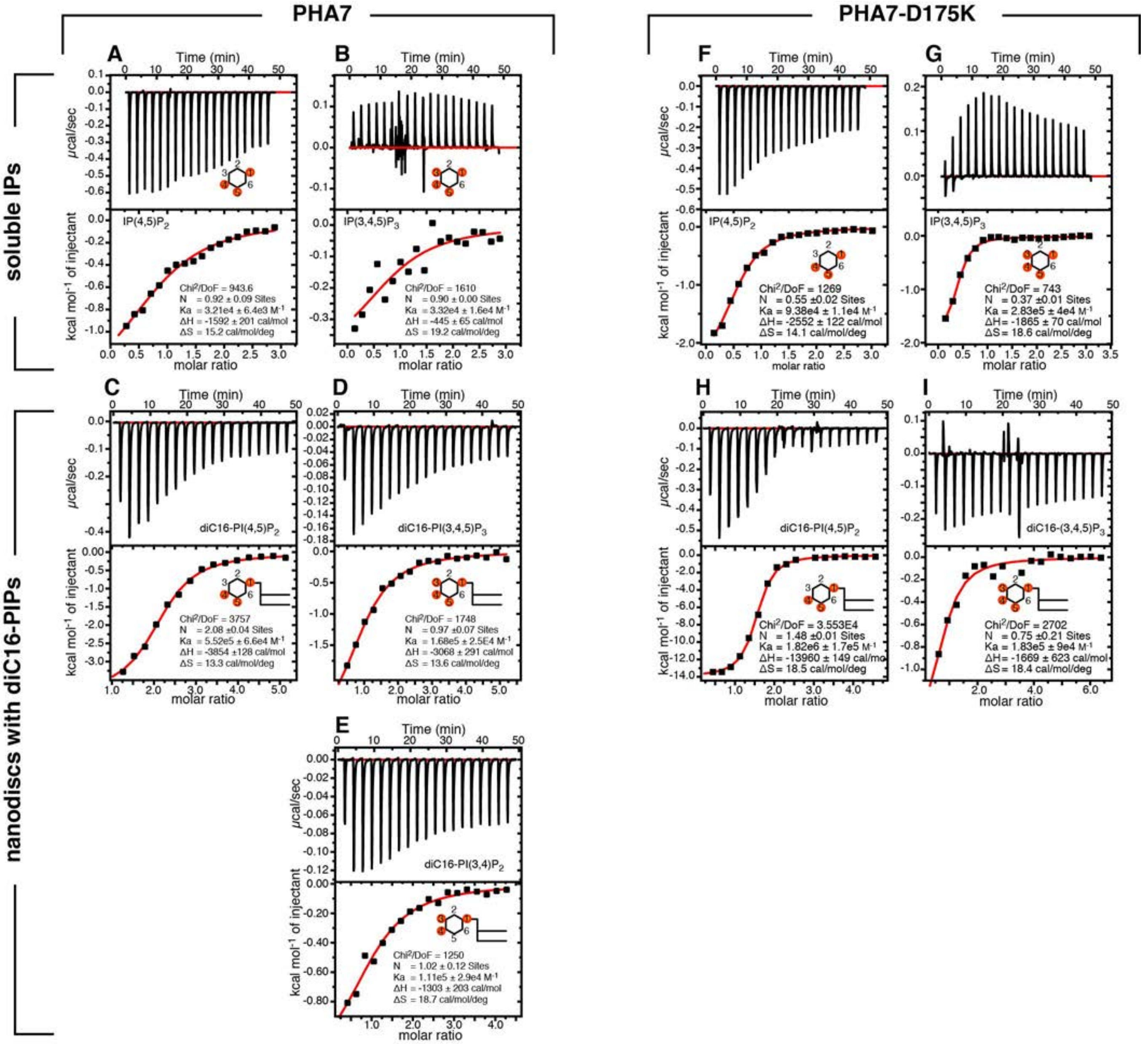

**Figure S4. Representative ITC binding isotherms** for titrations of wild-type PHA7 or mutant PHA7-D175K with soluble IPs (A, B, F, G) or nanodiscs containing 10% molar diC16-PIPs (C-E, H, I). Continuous red lines are the best fits of the data to a single-site binding model, used to extract the values of the dissociation constant ($K_d$).

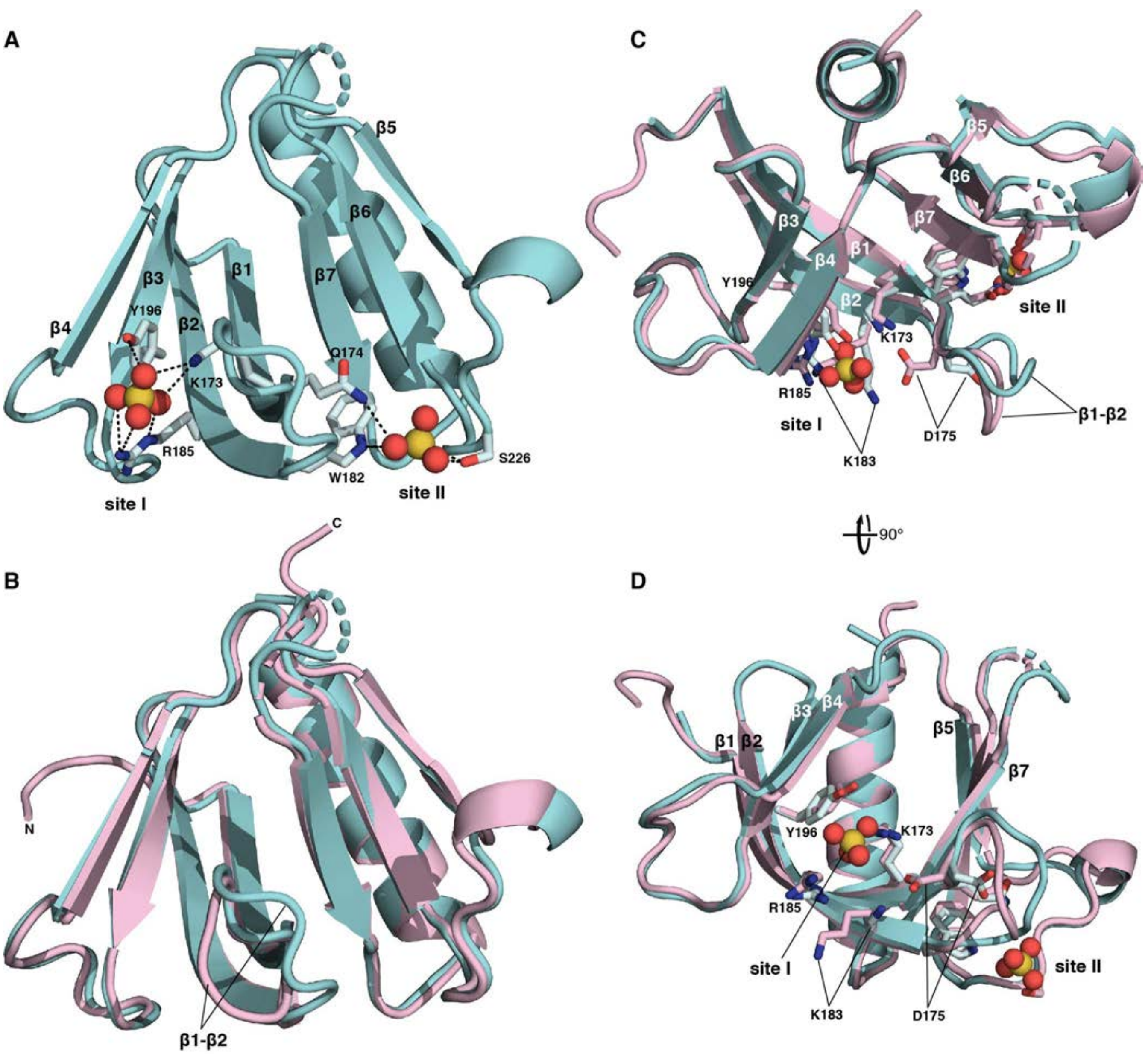


**Figure S5. Comparison of the structures of PHA7$_{APO}$ (pink) and PHA7$_{S}$ (cyan).** Sidechains involved in sulfate coordination are shown as sticks. Sulfate anions are shown as spheres (sulfur in yellow; oxygen in red). **(A)** View of the two binding sites (**I** and **II**) for sulfate of PHA7$_{S}$. **(B)** Structural superposition of PHA7$_{APO}$ and PHA7$_{S}$. The β1-β2 loop differs in the two structures. **(C, D)** Orthogonal view of the superimposed structures of PHA7$_{APO}$ and PHA7$_{S}$.

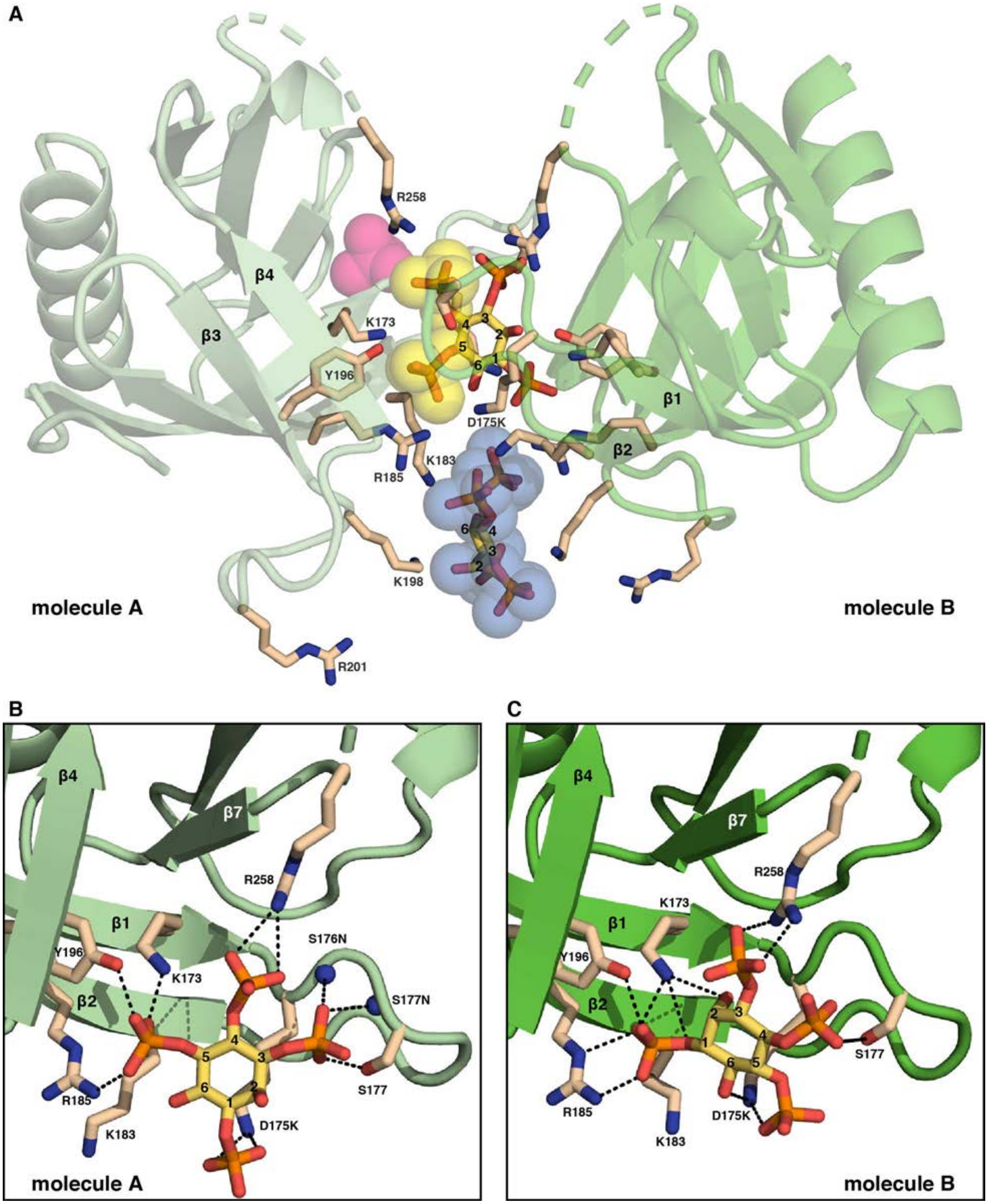


**Figure S6. Structure of PHA7-D175K. (A)** Two molecules of PHA7-D175K (pale green, dark green) in the crystallographic asymmetric unit coordinate two I(1,3,4,5)$P_4$ molecules sandwiched between them. One IP(3,4,5)$P_3$ molecule occupies the principal binding site. The fifth phosphate group on the inositol ring occupies binding site I and coincides with the sulfate anion that binds in the structure of PHA7$_S$ (cyan). The second IP(3,4,5)$P_3$ molecule binds more peripherally. **(B, C)** Association of IP(3,4,5)$P_3$ with site I in each of the two PHA7-D175K molecules of the crystallographic asymmetric unit.

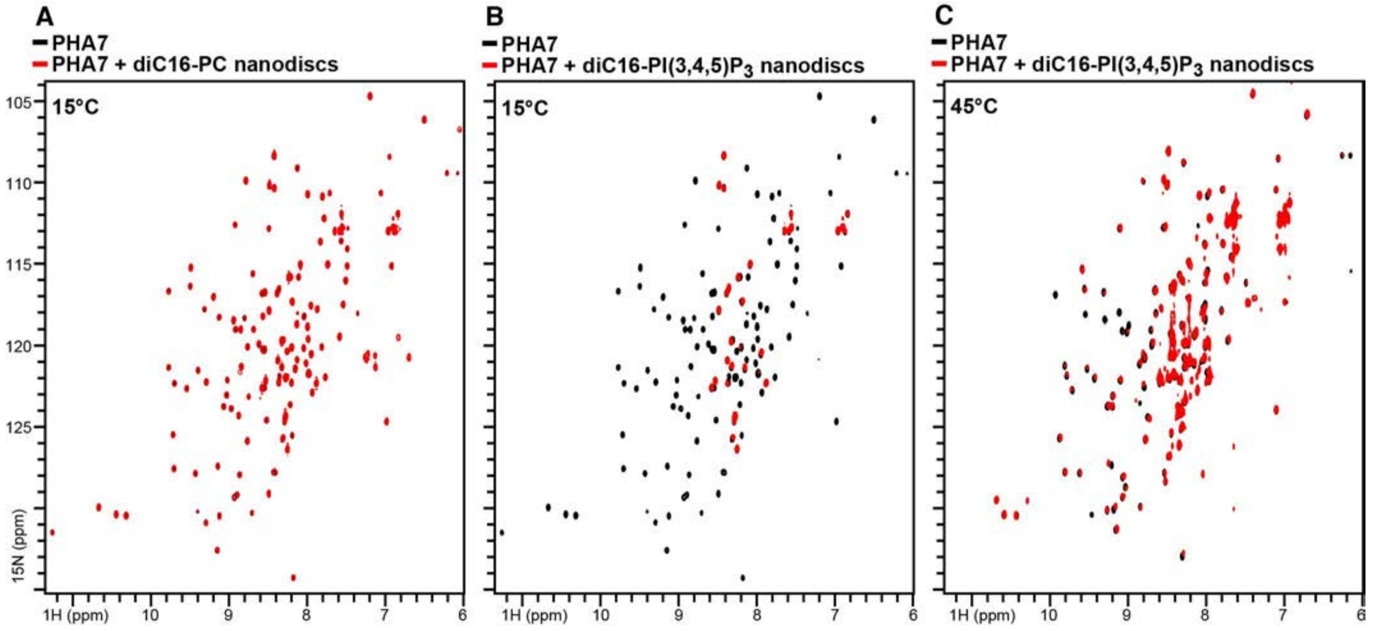


**Figure S7. Interaction of PHA7 with PIP nanodiscs.** $^{1}H/^{15}N$ spectra NMR of $^{15}N$-labeled PHA7 were obtained before (black) or after (red) addition of 1.5 molar equivalents of lipid nanodisc, containing: **(A)** 100% diC16-PC or **(B, C)** 10% diC16-PI(3,4,5)$P_3$. The spectra were acquired at 15°C (A, B) or 45°C (C).

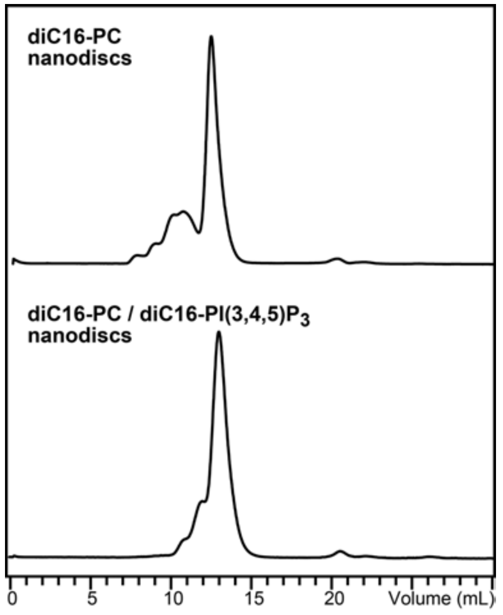


**Figure S8.** Size exclusion chromatography elution profile of nanodiscs used in this study. The major peak was collected for NMR experiments.

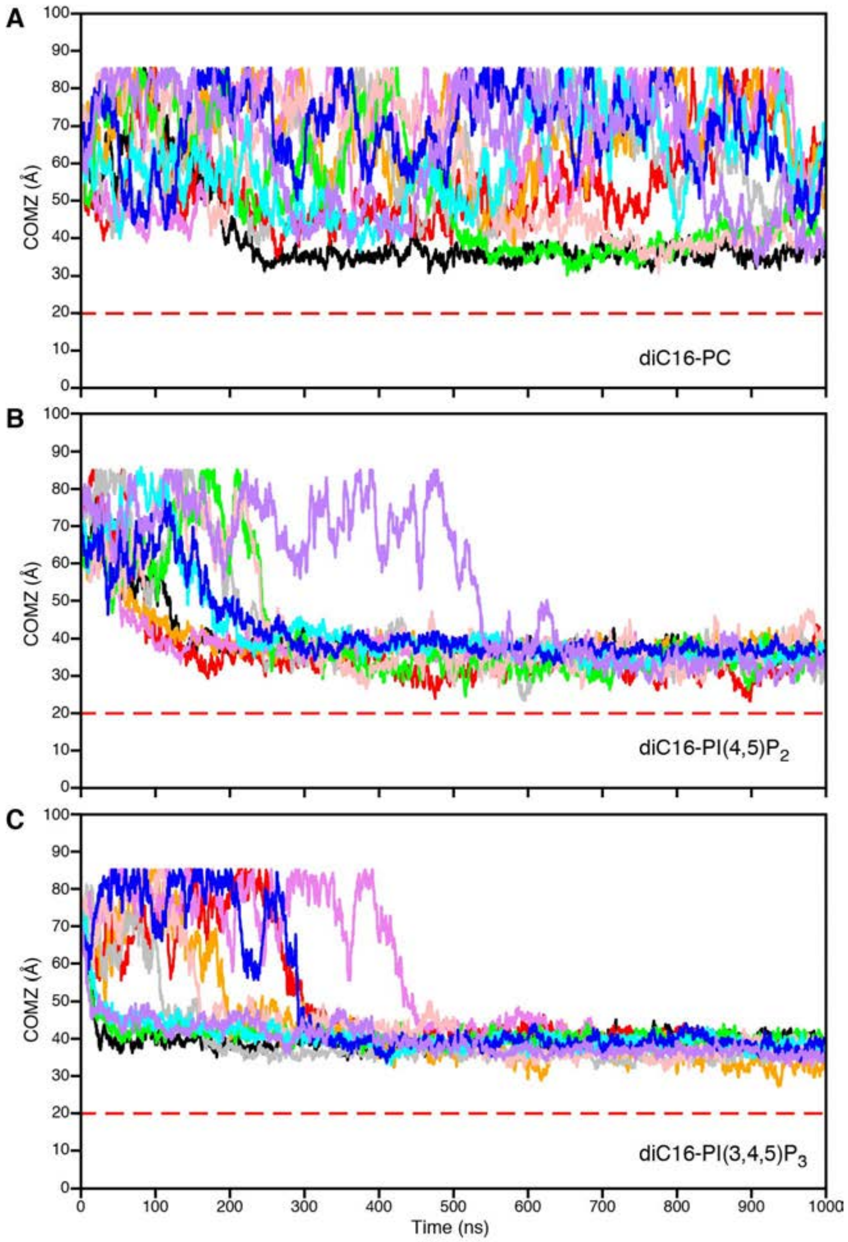


**Figure S9. MD simulations time series of the average distance of the center of mass of PHA7 from the center of the lipid bilayer membrane.** The distance is measured parallel to the lipid bilayer normal (Z axis) with the lipid bilayer center at Z = 0 Å. The average location of the lipid head groups is at Z = 20 Å (dashed red line). Different colors represent each of 10 independent simulations of PHA7 in: (A) pure diC16-PC membranes; (B) membranes containing 10% diC16-PI(4,5)$P_2$; or (C) membranes containing 10% diC16-PI(3,4,5)$P_3$. Through the periodic boundary conditions, PHA7 can associate with the lower bilayer leaflet (near Z = 85 Å). While PHA7 shows no specific binding to PC-only membranes, simulations with either PI(4,5)$P_2$ or PI(3,4,5)$P_3$ resulted in protein association with the membrane surface.

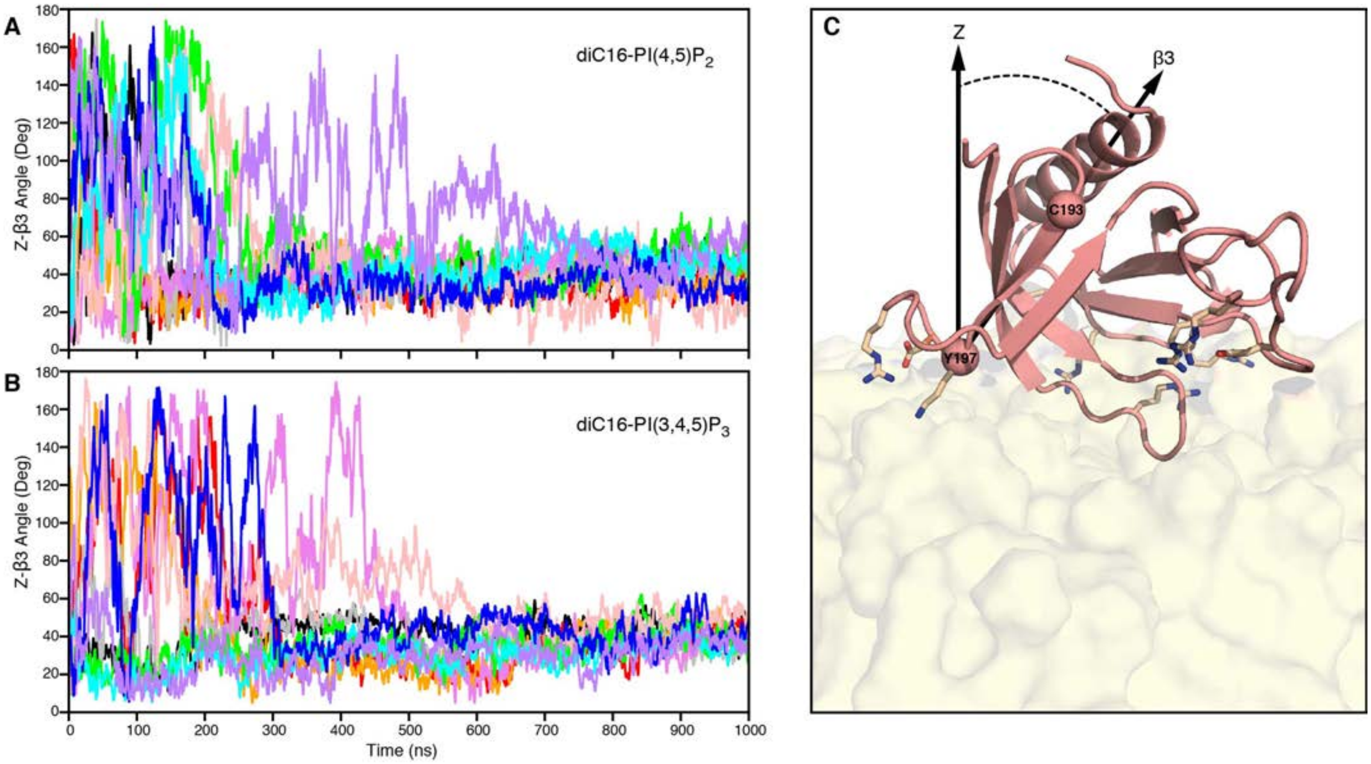


**Figure S10. MD simulations time series of the average preferred orientation of PHA7 on PIP membranes.** Different colors represent each of 10 independent simulations. **(A)** Membranes containing 10% diC16-PI(4,5)$P_2$. **(B)** Membranes containing 10% diC16-PI(3,4,5)$P_3$. **(C)** PHA7 protein orientation is represented by the angle between the membrane normal (Z) and the vector intersecting the CA atoms of C193 and Y197 (spheres) that define the β3 strand axis. The membrane is represented as a molecular surface (yellow). Key side chains with close (< 4 Å) contacts to PIP headgroups are shown as sticks.

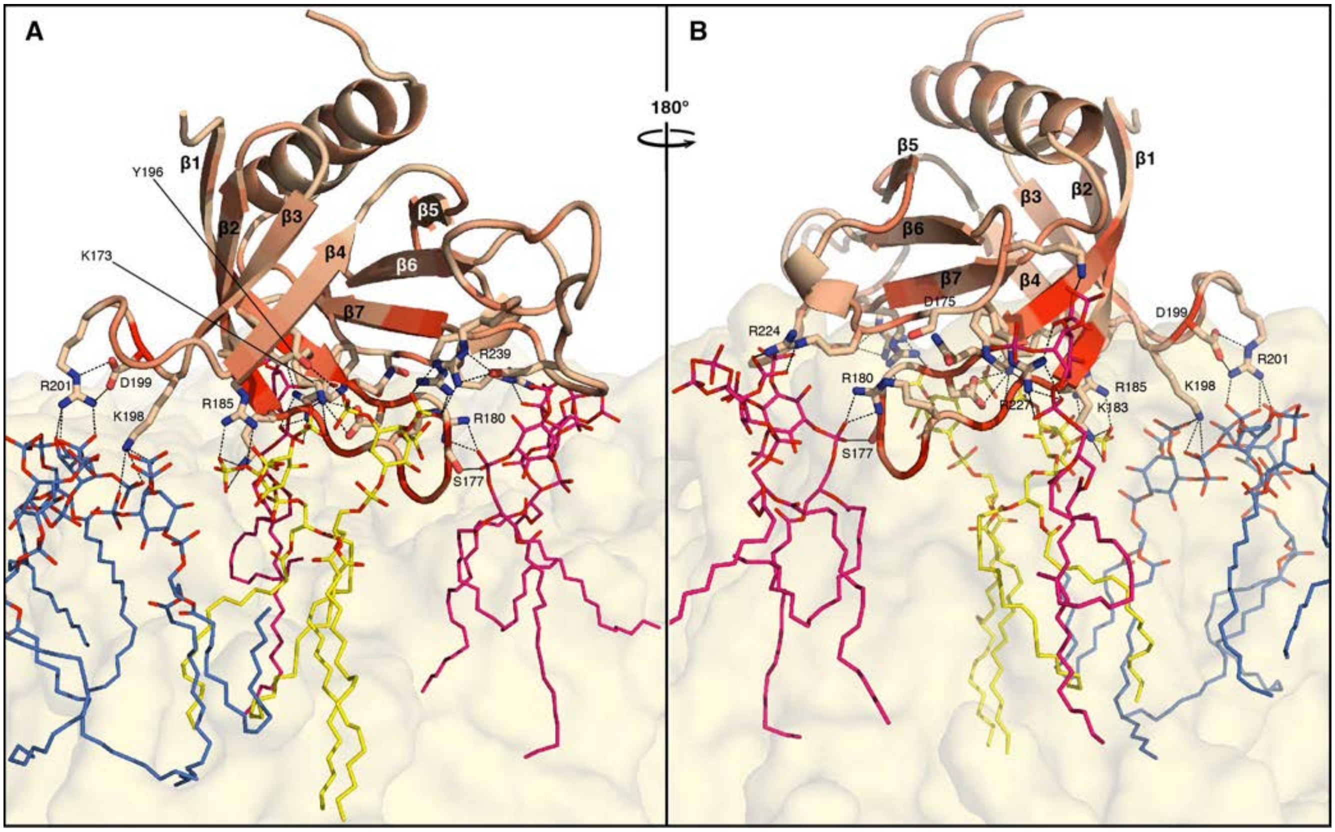


**Figure S11. PIP clustering induced by PHA7 association with the membrane surface. (A, B)** Snapshot (taken at 890 ns) of MD simulation of PHA7 with a diC16-PI(3,4,5)$P_3$ membrane. PHA7 colors reflect the magnitude of NMR chemical shift perturbation from 0 ppm (wheat) to the maximum value (red). Key side chains with close (< 4 Å) contacts to PIP headgroups are shown as sticks. The membrane is represented as a molecular surface (yellow). Colors of bound PIP molecules (lines) represent their association with binding sites **I** (yellow), **II** (pink), or **III** (blue).

**Table S1. Data collection and refinement statistics (molecular replacement) [a, b].**

| | **PHA7$_{APO}$ (PDB: 7KK7)** | **PHA7$_{S}$ PDB: 7KJO)** | **PHA7-D175K (PDB: 7KJZ)** |
|---|---|---|---|
| Data collection | | | |
| Space group | P $2_1$ $2_1$ $2_1$ | P $4_1$ | P $3_2$ 2 1 |
| Cell dimensions | | | |
| *a*, *b*, *c* (Å) | 57.31 59.06 78.47 | 64.83 64.83 59.23 | 77.47 77.47 82.56 |
| $\alpha$, $\beta$, $\gamma$ (°) | 90 90 90 | 90 90 90 | 90 90 120 |
| Resolution (Å) (outer shell) | 47.2 -2.80 (2.95 -2.80) | 43.7-1.45 (1.48- | 52.1-2.43 (2.53- |
| No. reflections * | 6980 (1000) | 1.45) | 2.43) |
| Wavelength (Å) | 1.116 | 46929 (4283) | 11134 (1136) |
| $R_{merge}$ | 0.071 (0.81) | 1.116 | 1.54 |
| $I / \sigma I$ | 16.0 (2.5) | 0.050 (0.362) | 0.058 (0.56) |
| $CC_{1/2}$ | 0.99 (0.89) | 22.2 (2.30) | 21.8 (3.9) |
| Completeness (%) | 100.0 (100.0) | n/d | 0.99 (0.91) |
| Redundancy | 6.8 (7.1) | 99.2 (99.0) | 99.9 (99.9) |
| | | 19.0 (18.1) | 10.4 (9.9) |
| Refinement | | | |
| Resolution (Å) | 47.2 – 2.80 | 45.84 –1.45 | 38.8 – 2.43 |
| No. reflections/test set | 6595 / 453 | 43487 / 2187 | 10443 / 668 |
| $R_{work}$ / $R_{free}$ | 0.212 / 0.276 | 0.155 / 0.171 | 0.201 / 0.236 |
| No. atoms | | | |
| Overall | 1701 | 1940 | 1794 |
| Protein | 1671 | 1695 | 1672 |
| Ligand/ion | 22 | 38 | 66 |
| Water | 8 | 207 | 60 |
| *B*-factors | | | |
| Overall | 100.0 | 29.3 | 71.3 |
| Protein | 100.0 | 27.4 | 70.2 |
| Ligand/ion | 106.1 | 51.9 | 112.5 |
| Water | 75.8 | 39.9 | 56.9 |
| R.M.S. deviations | | | |
| Bond lengths (Å) | 0.011 | 0.023 | 0.013 |
| Bond angles (deg) | 1.78 | 1.78 | 1.72 |
| Ramachandran favored/outliers (%) | 96.9 / 0.5 | 97.5 / 0.0 | 97.4 / 0.0 |

[a] Each data set was collected from a single crystal.
[b] Values in parentheses are for the highest-resolution shell.

**Table S2. Lipid compositions of each system generated for MD simulations.**

| **Membrane system** | **Number of lipid molecules per membrane system** | | |
|---|---|---|---|
| | **diC16-PC** | **diC16-PI(4,5)$P_2$** | **diC16-PI(3,4,5)$P_3$** |
| PHA7 + PC | 120 | 0 | 0 |
| PHA7 + PI(4,5)$P_2$ | 108 | 12 | 0 |
| PHA7 + diC16-PI(3,4,5)$P_3$ | 108 | 0 | 12 |
